# Inequality, heterogeneity, and chance: Multiple factors and their interactions

**DOI:** 10.1101/2024.05.31.596783

**Authors:** Hal Caswell, Silke F. van Daalen

**Affiliations:** Institute for Biodiversity and Ecosystem Dynamics, University of Amsterdam, Amsterdam, The Netherlands; Biology Department, Woods Hole Oceanographic Institution, Woods Hole, MA, USA

## Abstract

A heterogeneous population is a mixture of groups differing in vital rates. In such a population, some of the variance in demographic outcomes (e.g., longevity, lifetime reproduction) is due to heterogeneity and some is the result of stochastic demographic processes. Many studies have partitioned variance into its between-group and within-group components, but have focused on single factors. Especially for longevity, variance due to stochasticity is far greater than that due to heterogeneity. Here we extend analysis to multiple-factor studies, making it possible to calculate the contributions to variance of each factor and each of the interactions among factors. We treat the population as a mixture and use the marginal mixing distributions to compute variance components. Examples are presented: longevity as a function of sex, race, and U.S. state of residence, lifetime reproduction among set of developed countries and as a result of resource availability and pesticide exposure.

## 1 Introduction

A heterogeneous population is a mixture, made up of groups^1^ of individuals that differ in the demographic rates to which they are subject (Figure 1). Each group is characterized by a mean and a variance of some demographic outcome. That heterogeneity among individuals contributes to the variance, also among individuals, at the population level. Longevity is one example of a demographic outcome. Variance in longevity is of interest to demographers as a form of inequality (e.g., Vaupel, 1988; Edwards and Tuljapurkar, 2005; Vaupel et al., 2011; van Raalte et al., 2018; Permanyer and Scholl, 2019; Permanyer et al., 2023). Longevity can be generalised to include healthy longevity (with health defined in many ways) or occupancy of medical, infection, or other kinds of states (Caswell and Zarulli, 2018; Caswell and van Daalen, 2021). Variance in longevity has implications for health systems, pensions, estate planning, etc.

**Figure 1:**
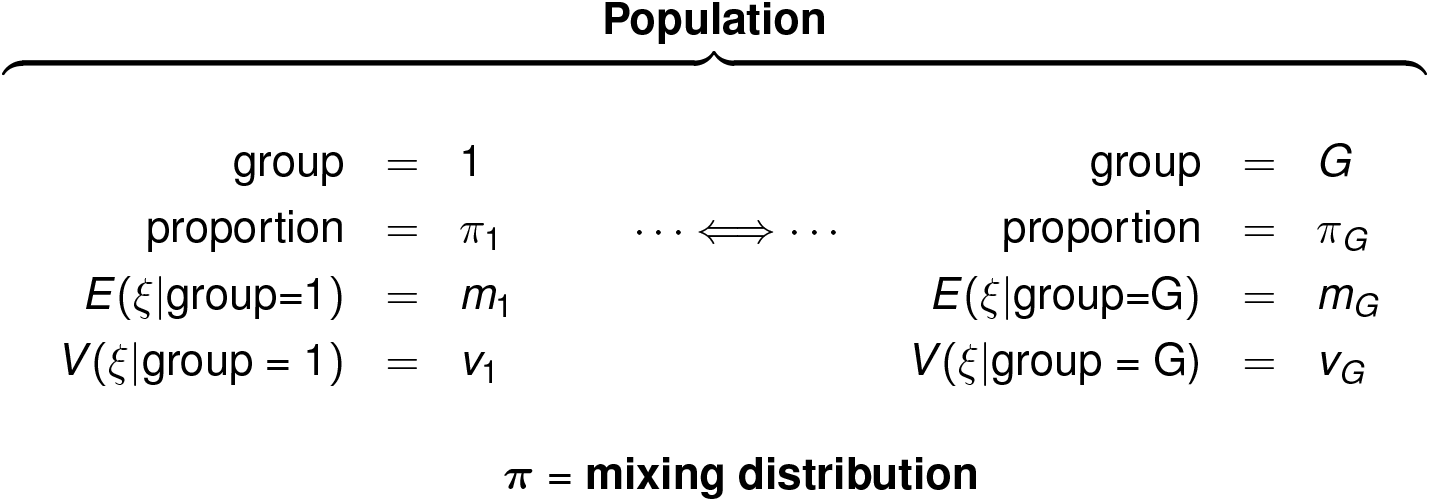
Diagrammatic representation of a heterogeneous population as a mixture of groups, each with its group-specific mean and variance for some demographic outcome ξ.

Lifetime reproduction, the number of offspring produced by a female over her lifetime, is an outcome of interest to evolutionary and anthropological demographers as a measure of the opportunity for selection (Crow, 1958; van Daalen and Caswell, 2024). It is often reported for hunter-gatherer populations (Hill and Hurtado, 1996; Blurton-Jones, 2016; Brown et al., 2009) or historical populations (Moorad et al., 2011; Courtiol et al., 2012).

Because demographic rates contain probabilities, demographic outcomes are random variables characterised by a distribution and a set of moments and other statistics. Because a heterogeneous population is a mixture, the distribution of the outcome is a *mixture distribution*, characterised by the moments of each group and the distribution of individuals among the groups. The latter distribution is called the *mixing distribution*.

### 1.1 Variance and variance decomposition

The variance in outcome in a heterogeneous population reflects both stochasticity within each group and heterogeneity among groups. The within-group variance is strictly due to chance in any outcome calculated from a life table, a Markov chain, or some equivalent machinery. Individual stochasticity is the only source for such variance, because the calculation explicitly applies the same probabilities to every individual at every age (Caswell, 2023).

The relative magnitude of the contributions from stochasticity and heterogeneity depend on many factors, including the type and magnitude of the heterogeneity. Thus it is essential to partition the variance into within-group and between-group components. Variances are often called “inequalities,” although that identification is more subtle than is usually appreciated (Caswell, 2023). In economic terms (e.g., Atkinson, 2015), variance due to heterogeneity corresponds to inequality of opportunity. Individuals in, e.g., different income groups have different opportunities to live a long life. Variance due to stochasticity corresponds to inequality of outcome: individuals in the same group differing in longevity because of the random outcomes of survival, even though they experience the same risks. Therborn (2014) provides the only statement, to our knowledge, of criteria that distinguish the kinds of heterogeneity that qualify as ‘inequality’.

The variance partitioning problem is classical. The variance in outcome among a heterogeneous set of individuals can be partitioned into between-group and within-group components using basic results from conditional probability. Consider a heterogeneous population that is a mixture of groups, the relative abundances of which is given by a mixing distribution ***π***. Let *ξ* denote some outcome; the variance in *ξ* is

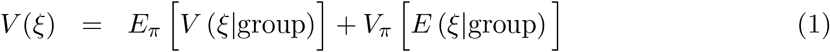

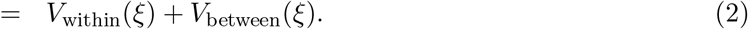

The within-group variance *V*_within_ is the expectation of the variance within each group, weighted by the distribution ***π***. The between-group variance *V*_between_ is the variance among the group means, again weighted by the distribution ***π***. This variance decomposition is a standard result in conditional probability (e.g., Rényi, 1970; Frühwirth-Schnatter, 2006) and in analysis of variance (ANOVA) (e.g., Kempthorne, 1957); indeed, the decomposition was introduced, along with the term ‘variance’ itself, by Fisher (1918).

### 1.2 Demographic outcomes: longevity and lifetime reproductive output

We will show results in this paper for variance decomposition for longevity and lifetime reproductive output (LRO). The means and variances (and other moments) of longevity are readily calculated by expressing the demographic rates in terms of absorbing Markov chains (pioneered by Feichtinger 1971; see Caswell (2001, 2006, 2009)). It is now known that, as a general rule, even extreme differences among groups, including those created by important socioeconomic variables, account for only a small fraction of the variance in longevity (see the overview in Caswell 2023).

The means, variances (and other moments) of LRO can be calculated using Markov chains with rewards (Caswell, 2011; van Daalen and Caswell, 2017). Stochasticity in LRO reflects both survival (which determines how long a woman has to reproduce) and fertility (which determines whether she reproduces or not at each age). These two components can be separated, and in developed, low mortality, countries the variance in LRO is increasingly accounted for by stochasticity in fertility (van Daalen and Caswell, 2015). But the contributions of heterogeneity and stochasticity to the variance in LRO are not yet well characterised. In one case, where the groups are defined by maternal age groups in a laboratory population, the variance components depend strongly on the environmental conditions (van Daalen et al., 2022). We will show some examples below, in Section 5.2.

### 1.3 Beyond the limitation to one factor

Studies of the variance in longevity and in lifetime reproduction usually define heterogeneity in terms of a single factor. When studies examine multiple factors, they are usually treated one at a time. Individuals, however, are heterogeneous in multiple factors operating simultaneously, and the variance is affected by all those factors and their interactions. The inability to evaluate the contributions of interactions is a major limitation to the study of heterogeneity and our goal here is to show how to analyse factorial studies, in which individuals are heterogeneous in two or more factors. We will present some two-factor and three-factor examples for both longevity and lifetime reproduction. There is no limitation to the number of factors.

## 2 Variance components

### 2.1 Notation

The following notation is used throughout this paper. Matrices are denoted by upper case bold characters (e.g., **U**) and vectors by lower case bold characters (e.g., **a**). Vectors are column vectors by default; **x**^T^ is the transpose of **x**. The vector **1** is a vector of ones, and the matrix **I** is the identity matrix. When necessary, subscripts are used to denote the size of a vector or matrix; e.g., **I**_*ω*_ is an identity matrix of size *ω* × *ω*. Matrices and vectors with a tilde (e.g., **Ũ** or **ã**) are block-structured; in this paper, blocks correspond to different factors. The notation ∥**x**∥ denotes the 1-norm of **x**. The symbol ⊗ denotes the Kronecker product. The vec operator stacks the columns of a *m* × *n* matrix into a *mn* × 1 column vector. When applied to an array with more than 2 dimensions, it stacks columns from all dimensions. We will make use of a `reshape` operator that is the inverse of the vec operator, changing the vector back into the array; see equation (17). On occasion, Matlab notation will be used to refer to the entries, rows, and columns of matrices. For example, **F**(*i*, *j*) is the (*i*, *j*) entry of **F**, and **F**(*i*, :) and **F**(:, *j*) refer to the *i*th row and *j*th column of the matrix.

Element-by-element operations apply to arrays of any dimension. The symbol ° denotes the Hadamard, or element-by-element product (implemented by .* in Matlab and by * in R). The symbol ⊘ is used to denote the Hadamard, or element-by-element quotient. Thus, for two matrices **A** and **B**,

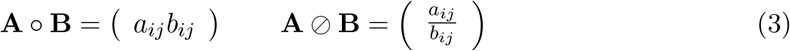

with the obvious restrictions that the objects must have the same dimension and, for the quotient, none of the entries of **B** can be zero.

### 2.2 Weighted means and variances

Let **x** be a vector of numbers and ***π*** a probability vector of the same length. The mean and variance of the entries of **x**, over the mixing distribution ***π*** are, in matrix notation,

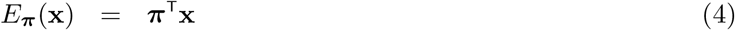

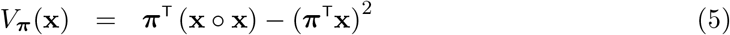

The second of these terms, the variance of *X* over a mixing distribution, will appear so commonly that we define it as a function 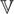 (**x, *π***) that returns the variance, over the distribution ***π***, of the vector **x**:

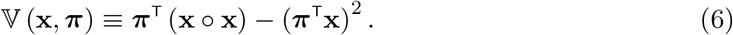

### 2.3 One-factor designs

Let us review the simplest case: a one-factor design in which a population (or some other set of individuals) is divided into groups based on a single factor which we call A (e.g., income), with levels 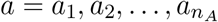. The population is a mixture, and individuals are distributed among the groups according to a mixing distribution π. Let *ξ* be the demographic outcome of interest. The vectors of means and variances are

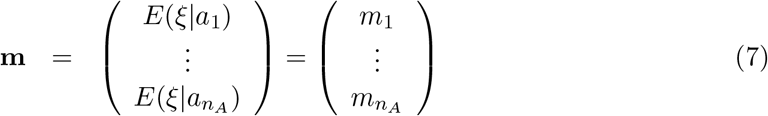

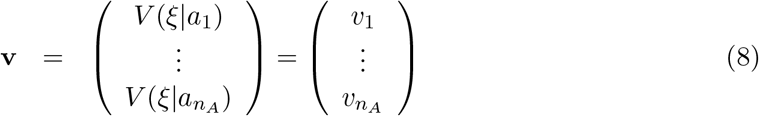

In terms of these vectors, the variance decomposition in equation (2) becomes

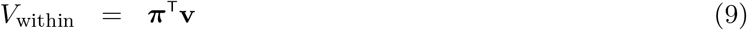

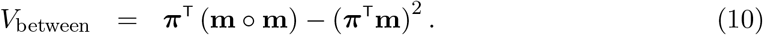

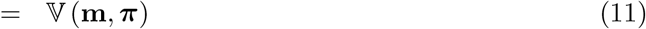

The relative contribution of the within- and between-group components to *V*(*ξ*) is measured by

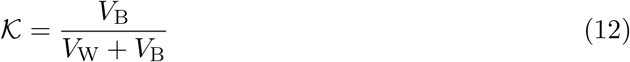

which measures the fraction of the total variance due to heterogeneity among groups; it is referred to as the intraclass correlation coefficient in quantitative genetics (Falconer, 1960). Its square root is called the correlation ratio in probability theory (Rényi, 1970).

This analysis, for one-way designs, is straightforward to calculate (given the use of the multistate demographic model that permits inclusion of groups and calculation of variances) and has now been used repeatedly. It has resulted in a series of quantitative measurements of the contributions of heterogeneity (among groups) and individual stochasticity (with groups) to variance in:

- longevity among frailty classes in human populations (Hartemink et al., 2017),
- longevity among latent mortality groups in laboratory populations of insects (Hartemink and Caswell, 2018; van Daalen and Caswell, 2020) and in a wild seabird population (Jenouvrier et al., 2018),
- longevity among human populations in Scotland, characterised by a neighbourhood deprivation index (Seaman et al., 2019),
- lifetime reproductive output in an herbaceous plant, with groups defined by the fire status of the local environment (van Daalen and Caswell, 2020),
- longevity and lifetime reproductive output in laboratory populations of a rotifer, with groups defined by maternal age (van Daalen et al., 2022),
- longevity and lifetime reproductive output among many populations of plants and of animals, with groups defined by species and/or populations (Varas Enríquez et al., 2022; Caswell, 2023), adult longevity in a tephritid fruit fly, with groups defined by larval diet (Caswell, 2023),
- longevity in human populations with groups defined by income, education, or occupation (Caswell, 2023), and healthy longevity in human populations, with groups defined by European country of residence (Caswell, 2023).

## 3 Factorial designs: two factors

The multi-factor analysis is based on multi-way tables of the means, variances, and mixing distribution. The calculation of variance components uses marginal distributions calculated from those tables. Consider two factors, labelled A and B (e.g., sex and race) with levels 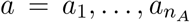 and 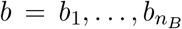. For example, for sex *n_A_* might be 2, and for race *n_B_* could be two, or five, or some other number depending on the information collected in vital statistics registries. The means, variances, and mixing distributions are defined in the 2-dimensional arrays

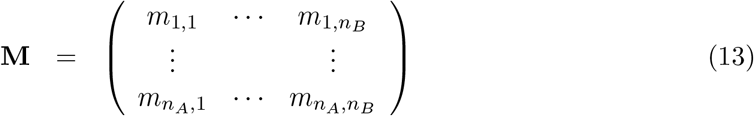

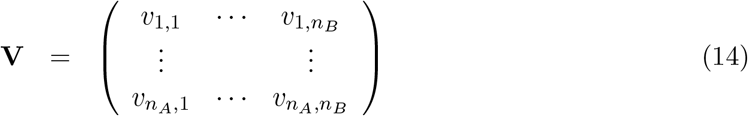

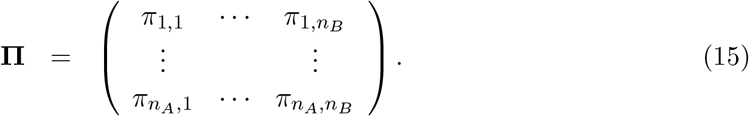

Each of these arrays is of dimension *n_A_* × *n_B_*. The mixing distribution array satisfies

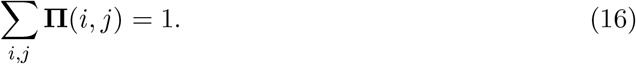

The vec operator applied to any of these arrays produces a vector, of dimension *n_A_n_B_* ×1, by stacking the columns of the array. The inverse of the vec operator is the `reshape` operator, which takes as its arguments a vector and a pair of dimensions, and produces an array of those dimensions; thus

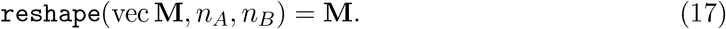

### 3.1 Within-group and between-group variances

Treating each of the *n_A_n_B_* combination of factors A and B as a group, the within- and between-group variance components are calculated from the entries of **M, V**, and **Π**. The within-group variance is the mean, over the mixing distribution, of the variances in each group,

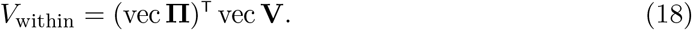

The between-group variance is given by applying equation (6) to the means of all the factor combinations:

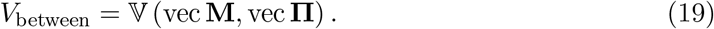

This between-group variance measures the overall contribution of heterogeneity, over all the groups, of all the factors, with individuals distributed according to **Π**. It makes no distinction between the contributions of A, B, or the AB interaction, but if the goal is a measure of how heterogeneity in factors contributes to inequality of outcomes, *V*_between_ is the answer.

### 3.2 Partitioning the between-group variance into factor effects

In the two-factor case, *V*_between_ is due to contributions from each factor (*V*_A_, *V*_B_) and their interaction (V_AB_). These components are calculated from the marginal means and marginal mixing distributions corresponding to each factor and the interaction. Use subscripts to identify the marginal means (e.g., **m**_*A*_ as the vector of means for each level of A, marginalising over levels of B), and let a, b denote levels of A and B respectively. The array of marginal means for factor A is obtained by calculating the average, weighted by the mixing distribution, over the levels of factor B. Recalling the definitions of the Hadamard product and Hadamard quotient in Section 2.1, we have

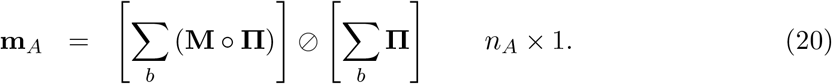

The same pattern holds for the marginal mean for B:

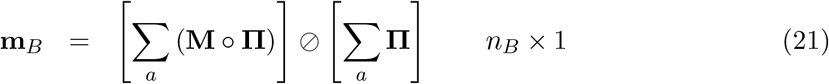

and for the marginal mean for AB

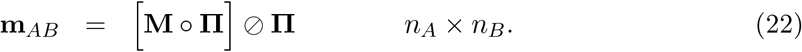

The marginal mean for factor A is obtained by summing over levels of B; that for factor B is obtained by summing over levels of A. The marginal mean for the combination AB is not really marginal; there are no other factors over which to sum. The marginal means for factors A and B are one-dimensional vectors. The marginal mean for the AB interaction is a two-dimensional array (it is, in fact, just the array **M**).

The marginal mixing distributions are obtained from the array **Π**,

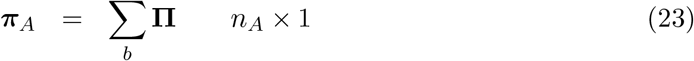

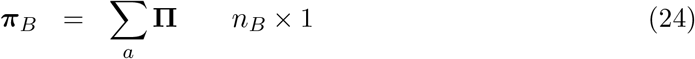

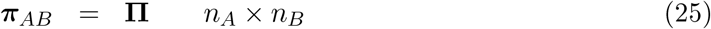

Matlab makes these calculations easy to implement; the command corresponding to equation (20), for example, is

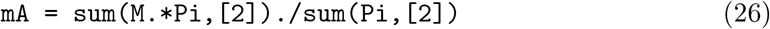

Finally, the variance components due to factors A and B and the interaction AB are calculated by applying the function 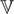 (·, ·) defined in equation (6) to the marginal means and marginal mixing distributions:

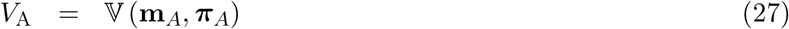

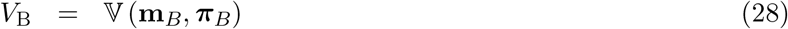

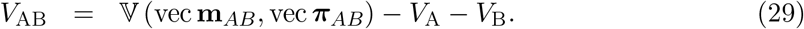

The marginal mean array **m**_*AB*_ contains the effects of A and B as well as the interaction; thus the interaction variance *V*_AB_ is obtained by subtracting *V*_A_ and *V*_B_ from the variance among all factor combinations.

### 3.3 Requirements for the factorial mixing distribution

The calculation of *V*_between_ treats vec **M** and vec **V** as vectors containing values for each of the groups. There is no restriction on the mixing distribution **Π** except, of course, that its entries sum to one.

But we want to partition **V**_between_ into its components, and to do this the mixing distribution **Π** requires some careful attention. Statistics texts are unanimous in stressing the importance of equal sample sizes in all treatment combinations in the analysis of variance (ANOVA) in a factorial experiment. The sample sizes play the role of the mixing distribution in our probability calculations. It has long been known (Yates, 1934) that unequal sample sizes make it impossible to calculate variance components. A recent text says

“When the sample sizes for each cell are unequal, the two-way analysis of variance for factor effects becomes complex. The component sums of squares in the analysis of variance are no longer orthogonal; that is, they do not sum to the total sum of squares. The least squares method for obtaining the best estimates of the parameters is rather complicated in the fixed effects model and the best analysis has not been and probably will not be found for the random effects models” (Sahai and Ageel, 2012).

There exist two classes of mixing distributions that permit calculation of variance components. One is the flat, or balanced distribution, which corresponds to equal sample sizes and assigns equal weight to all factor combinations. The other is the class of rank-one, or proportional mixing distributions (Yates, 1934; Kirk, 1982, and many others). Such a mixing distribution, when written as an array like equation (15), has proportional rows and proportional columns. The flat distribution is a special case of the rank-one distribution.

A rank-one mixing distribution array is one that can be assembled from its marginals. Let ***π***_*A*_ and ***π***_*B*_ be the marginal distributions among the levels of factors A, B. The mixing distribution array is

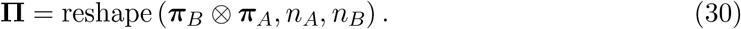

Every column is proportional to ***π***_*A*_ and every row is proportional to ***π***_*B*_. Note the order of the subscripts.

#### 3.3.1 The mixing distribution is a useful tool

Our goal is to understand the sources of variance in some outcome, within some population. But what population? The mixing distribution describes the structure of the population that we are interested in, and over which the variance is to be calculated, in terms of the proportions of the population in each of the heterogeneity groups. The proper question is not “what is the *correct* mixing distribution” but “what mixing distribution answers the question we are interested in?” A flat mixing distribution gives every group equal representation in studying the effects of heterogeneity.

It is a particularly powerful tool, not because many stage × group combinations have equal sample sizes,^2^ but for the same reason equal sample sizes are desirable in designed experiments. If you want to quantify the effects of some number of factors, it is wise to design your experiment with equal sample sizes in each treatment combination, because doing so maximises the ability to extract information obtained from the ranges of both variables. Thinking of a variance component calculation as a kind of numerical experiment to evaluate the effects of stages and groups is a valuable perspective.

Alternatively, the mixing distribution may reflect the relative population sizes of groups (subject to the proportional restriction). If some groups are much larger than others, the population variance will primarily reflect variance within those groups and pay no attention to the much smaller groups; we will see an example below.

## 4 Factorial designs: three factors

Variance partitioning for three factors follows the same pattern as that for two factors. We present the formulas briefly here; the extension to an arbitrary number of factors should be clear.

Consider three factors are labelled A, B, and C (e.g., sex, race, and state of residence), with levels 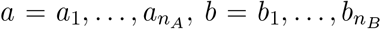, and 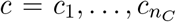. The arrays **M** of means, **V** of variances, and **Π** of mixing probabilities are now 3-dimensional. Again, the mixing distribution array satisfies

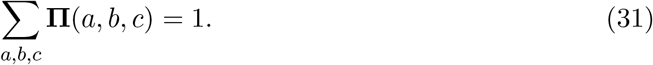

The vec operator produces a vector, of dimension *n_A_n_B_n_C_* × 1 by stacking the columns of the array in the order A, then B, and then C.

The mixing distribution array must be assembled from its marginals. Let ***π***_*A*_, ***π***_*B*_, ***π***_*C*_ be the marginal mixing distributions among the levels of factors A, B, and C, respectively. Then the rank-one mixing distribution array, generalizing that for two factors in (25) is

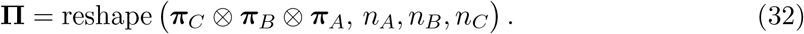

The extension to more than three factors follows the same logic.

### 4.1 Within- and between-group variances

As in Section 3, the within-group and between-group variances are calculated treating all *n*_*A*_*n*_*B*_*n*_*C*_ factor combinations as groups. Then, just as in the two-factor case,

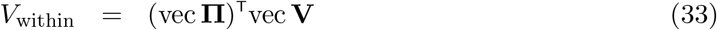

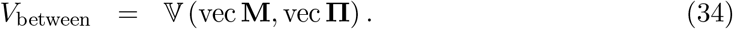

As in the two-factor case, *V*_between_ gives the contribution to variance of all the heterogeneity among groups, but no information on the contributions of the factors and their interactions.

### 4.2 Components of the between-group variance

The between-group variance *V*_between_ is partitioned into components due to the main effects of each factor (*V*_A_, *V*_B_, *V*_C_), the two-way interactions (*V*_AB_, *V*_AC_, *V*_BC_) between pairs of factors, and the three-way interaction (*V*_ABC_). These components are calculated from the marginal means and marginal mixing distributions corresponding to each factor and each interaction.

The array of marginal means for factor A is obtained by averaging over the dimensions other than those for factor A:

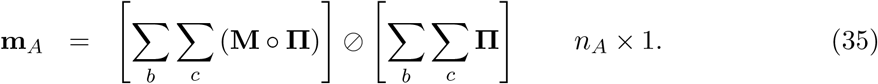

The same pattern holds for the other marginal means:

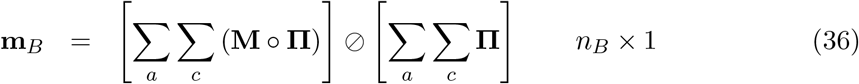

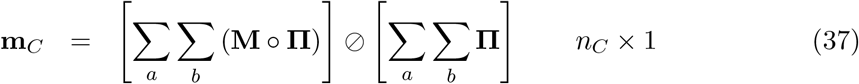

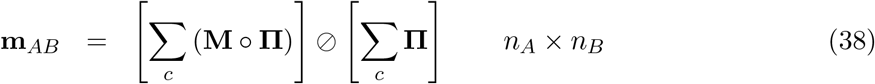

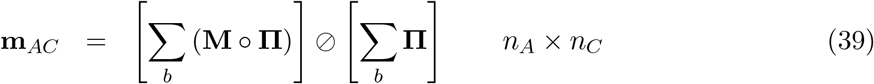

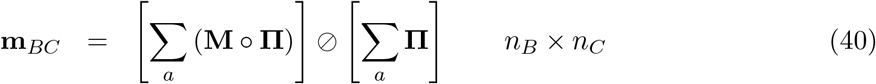

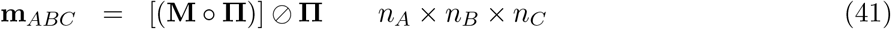

The marginal mixing distributions are obtained from the array **Π**,

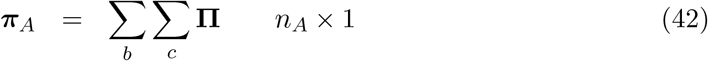

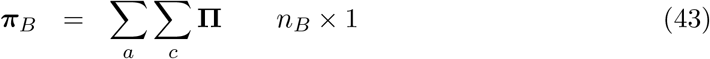

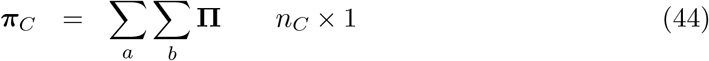

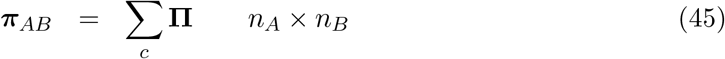

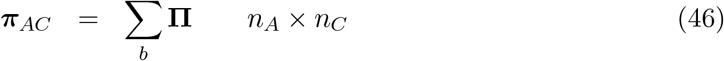

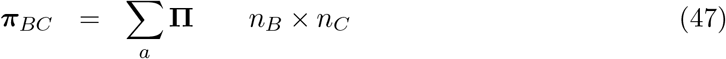

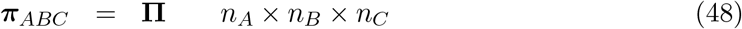

The calculations are readily expressed as Matlab commands; for example the command corresponding to equation (35), for example, is

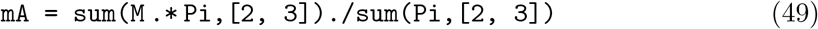

The second argument ([2,3]) in the sum command indicates the dimensions over which summation takes place.

The variance components due to each of the factors and interactions are calculated by applying the function 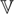 (·, ·) in equation (6) to the vectors obtained by applying the vec operator to the marginal mean arrays

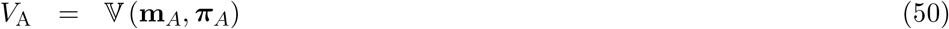

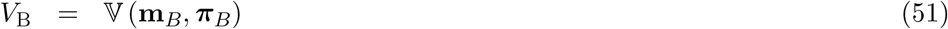

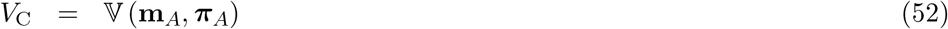

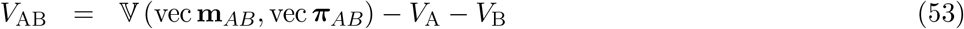

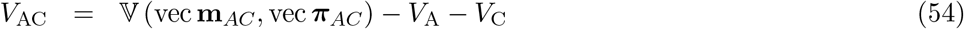

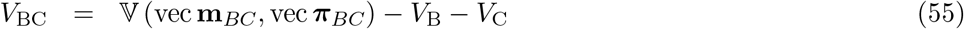

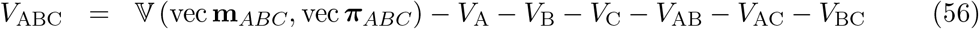

The arrays for the two-way interactions also include the one-factor effects, so the one-factor variances are subtracted to obtain the two-factor variances. The variance due to the three-factor interaction has the one-factor and two-factor variances subtracted.

### 4.3 The interpretation of interactions

The interpretation of interactions in factorial experiments has always been a challenge (e.g. Steel and Torrie, 1960; Sahai and Ageel, 2012). A large component of variance due to an AB interaction makes it difficult to say what the effects of A and B are, because the effect of A depends on the level of B, and vice-versa. The situation becomes even more difficult, of course, for three-way or higher interactions. If the contributions of interactions to variance are small, they can be ignored. Note that we have no operational definition of ‘small’ such as is provided in ANOVA by tests of the statistical significance of the interactions. However, the difficulty of interpreting interactions doesn’t change the fact that they substantively interesting. Knowing that two factors interact is an important finding and invites further study to understand how that interaction works.

If the factors have additive effects on *ξ*, the interaction variances are all zero (Caswell, unpublished results).

## 5 Examples

We present here several examples of multi-factor variance decompositions: two studies of longevity and two of lifetime reproductive output. At this point the results are intended to serve only as examples of choices of mixing distributions and of the kinds of results obtained. As always when a new analytical method is deployed, interpretation of the results is still developing.

### 5.1 Longevity and lifespan

Variance in longevity, often referred to as inequality or disparity in lifespan, has been analysed across a variety of social, economic, and biological variables. Partitioning of the variance into components has revealed that, even when group differences are very large, they contribute only a small fraction of the total variance (see an overview in Caswell 2023). Because longevity is the outcome of a lifetime of repeated probabilistic survival events, it is subject to a large degree of individual stochasticity.

However, these analyses have been limited to single factors. Here, we report two examples of multi-factor studies, one examining variance due to sex and race, the other examining sex, race, and state of residence. In these examples, we calculated the mean and variance of longevity using Markov chain methods (Feichtinger, 1971; Caswell, 2001, 2006), but they could equally have been calculated from a set of life tables.

#### 5.1.1 Variance in longevity: sex and race

Differences in longevity between males and females are well known; in almost every case, women live longer on average than men. Differences among racial and ethnic groups are also well known. Here we explore the variance in longevity for males and females in racial and ethnic categories in the United States. The 2020 life tables for the United States (Arias and Xu, 2022) classify individuals as male or female and into five racial and ethnic categories: Hispanic (H), Non-Hispanic American Indian and Alaska Native (NHAIAN), Non-Hispanic Asian (NHA), Non-Hispanic Black (NHB), and Non-Hispanic White (NHW). The arrays of mean longevity (life expectancy) and variance in longevity are

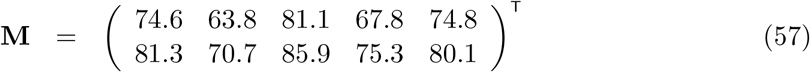

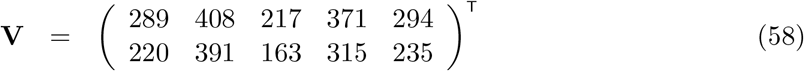

with males in the first row and females in the second row. These differences in life expectancy among ethnic groups and between the sexes are typical for those variables.

Two mixing distributions suggest themselves, asking different questions. A flat mixing distribution treats each sex-race combination equally in calculating its contribution to the variance. It provides information on how the difference in conditions experienced by these groups contributes to the variance among individuals in length of life, and thus tells something about the sex-race groups per se. However, the groups have quite different representation in the U.S. population. A mixing distribution proportional to the population sizes of each racial group, with sexes treated as equal, provides a decomposition of the variance in longevity among a hypothetical set of individuals selected at random from the population. It tells something about how the different conditions experienced by the groups *and* the population structure of those groups contribute to the variance in length of life.

Using population figures from 2020 (U.S. Census Bureau, Population Division, 2022), and setting the sexes equal, the marginal distributions for the flat mixing distribution are

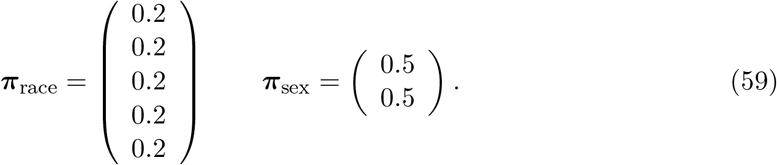

The marginal distributions for the population-weighted mixing distribution are

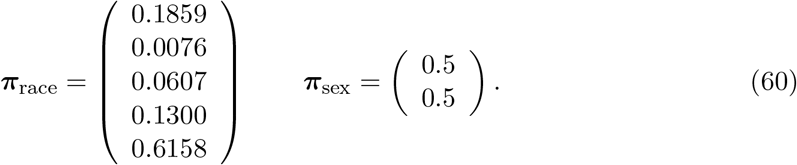

In the rank-one mixing distribution, the Hispanic and NH White groups account for 80% of the mixture, giving much less weight to the other racial categories. The rank-one mixing distribution is given by

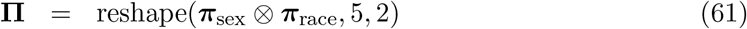

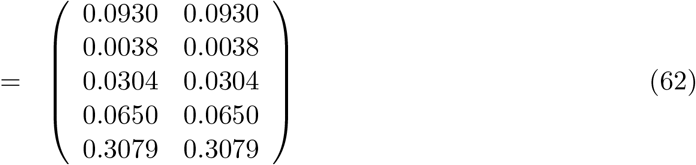

The variance decompositions obtained from the flat and the population-weighted mixing distributions are given in Table 1. Under the flat mixing distribution, the between-group variance accounts for 12.5% of the total variance. Racial differences contribute about 3 times the variance of sex differences, and the interaction contributes only a small fraction. When attention shifts to the population-weighted mixing distribution, the contribution of racial-ethnic heterogeneity shrinks from 31.5 to 7.4, as a result of the dominance of the population by the Hispanic and Non-Hispanic White groups. Combining both factors, heterogeneity accounts for only 5.6% of the variance.

**Table 1:** The components of variance due to race, sex, and their interaction, of the population of the United States, 2020, The rank-one mixing distribution is constructed by setting the marginal distribution of races proportional to their abundance in the U.S. population in 2020.

| Flat mixing |  | Population-weighted mixing |  |
| --- | --- | --- | --- |
| Component | Variance | Component | Variance |
| A=Race | 31.5 | A=Race | 7.4 |
| B=Sex | 9.7 | B=Sex | 8.5 |
| AB=Race × sex | 0.3 | AB=Race × sex | 0.2 |
| (between-group) | 41.5 | (between-group) | 16.1 |
| Stochasticity | 290.3 | Stochasticity | 269.3 |
| Total | 331.8 | Total | 285.4 |
| $\mathcal{K}$ | 0.125 | $\mathcal{K}$ | 0.056 |

#### 5.1.2 Variance in longevity: sex, race, and state of residence

In a country as large and diverse as the United States, there can be appreciable regional differences in mortality. The U.S. Census Bureau provides male and female life tables for each of the 50 states and the District of Columbia (Arias et al., 2022). The differences in life expectancy among states are large, and are partly a reflection of the political affiliations and policies of the states, with occupants of more liberal states experiencing longer life expectancies (Montez and Farina, 2021; Montez et al., 2020). An earlier study, at the level of counties in the U.S., found such large differences that the authors suggested the existence of “eight Americas” (Murray et al., 2006).

As an example of a three-factor analysis, we consider United States life tables by sex, race, and state of residence as given in (Wei et al., 2012). Unfortunately, the results are not directly comparable with the race×sex analyses in Section 5.1.1 because only two racial groups, White and Black, were reported in these data. Only 41 states were included, because the sample sizes for the Black population in the other states were considered too small to provide reliable estimates of mortality. However, we present it as an example of the potential for a three-factor interaction.

Table 2 shows the variance decomposition with a flat mixing distribution. The largest main effect variance component is that due to sex, followed by race, and then state of residence. The two-way interactions are small, as is the three-way interaction. Taken together, the interactions account for just over 5% of the between-group variance. Within-group stochasticity again accounts for most of the variance, and the variance ratio 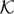 = 0.059. That is, the combined effects of the three factors and their interactions account for only 5.9% of the variance in longevity.

**Table 2:** Components of variance in longevity due to sex, race, and U.S. state of residence.

| Flat mixing |  |
| --- | --- |
| Component | Variance |
| A=sex | 9.23 |
| B=race | 7.50 |
| C=state | 1.37 |
| AB=sex × race | 0.061 |
| AC=sex × state | 0.209 |
| BC=race × state | 0.567 |
| ABC=sex × race × state | 0.170 |
| (between-groups) | 19.11 |
| Stochasticity | 303.04 |
| Total | 322.14 |
| $\mathcal{K}$ | 0.059 |

### 5.2 Variance in lifetime reproductive output

Lifetime reproductive output (LRO), sometimes called lifetime reproductive success (LRS), is the number of children produced by a female over her lifetime. The mean of LRO is the net reproductive rate R_0_. The mean LRO conditional on surviving through reproductive years is the total fertility rate TFR.

These mean values are widely used, but individuals are highly variable in their lifetime reproduction. LRO is a random variable determined by survival and reproduction, both of which are stochastic processes. The variance in lifetime reproduction is of equal interest and significance to the variance in longevity, but the idea of “reproductive inequality” corresponding to “lifespan inequality” appears to have drawn little attention.

The mean and variance (indeed, all the moments) of lifetime reproduction can be calculated using Markov chains with rewards (MCWR); see Caswell (2011); van Daalen and Caswell (2017, 2020) for details of the calculation. Briefly, a Markov chain describes individual development, survival, and transition among the possible states of the life history. The rewards are the offspring produced along the life history trajectory; the rewards are accumulated until death. The mean and variance of lifetime fertility are calculated from the demographic rates; they depend on the fate of the individual and on the outcomes of the chances of reproduction at each point in that lifetime.^3^

The crucial difference between LRO and TFR is that the former accounts for mortality of mothers or potential mothers in calculating the lifetime performance. LRO measures the amount of reproduction, and its variance, that is actually produced during the life of an individual. Extensive comparisons of the variance, skewness, and other statistics of LRO, for humans and other species, have been reported (Caswell, 2011; van Daalen and Caswell, 2015, 2024; Varas Enríquez et al., 2022) but few studies have partitioned variance in LRO into contributions from heterogeneity and stochasticity. van Daalen et al. (2022) analysed the contribution of heterogeneity in maternal age to variance in LRO in a rotifer. Snyder and Ellner (2018, 2016) use a different approach, but also find that the contribution of trait heterogeneity to the variance in LRO is small.

Here, we present two examples of multi-factor studies in which variance in lifetime reproduction is partitioned into contributions from heterogeneity and stochasticity. One is a comparison of lifetime fertility among human females in 31 developed countries over a 40-year time interval, treating country and time as factors. The other is a laboratory study of the effects of food limitation and pesticide exposure on a rotifer.

#### 5.2.1 Variance in reproduction over time and among countries

van Daalen and Caswell (2015) used Markov chains with rewards to explore the statistics of lifetime reproduction (mean, variance, skewness, standardised variance) in a set of developed countries during the second demographic transition, using data from the Human Mortality Database and the Human Fertility Database.

Strictly as an example of the calculations, we present an analysis of a two-factor variance decomposition, choosing as factors two years (1960 and 2000), and the 31 countries for which data were available for both years.^4^ We computed the within-group and between-group components of variance, comparing two mixing distributions. One is flat, treating each country-year combination as equally important. The other creates marginal mixing distributions proportional to population size, measured at age 0. This accounts for the distribution of population sizes of individuals beginning their ‘lifetime reproduction’ that would result from randomly selecting newborn females in proportion to their abundance.

Table 3 shows the components of variance between years, among countries, the interaction of countries and years, and the variance within country-year combinations due to stochasticity. With a flat mixing distribution, the variance due to years is more than twice that due to countries. The interaction of years and countries is very small. The within-factor variance due to stochasticity is large, and the variance ratio 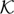 = 0.21. That is, only 21% of the variance in lifetime reproduction is due to the historical changes over this 40 year period and the political and social differences among countries over these two years. Almost 80% of the variance is due to stochasticity from the random outcome of survival and fertility. Within that 21% of the variance due to heterogeneity, 91% is due to main effects.

**Table 3:**
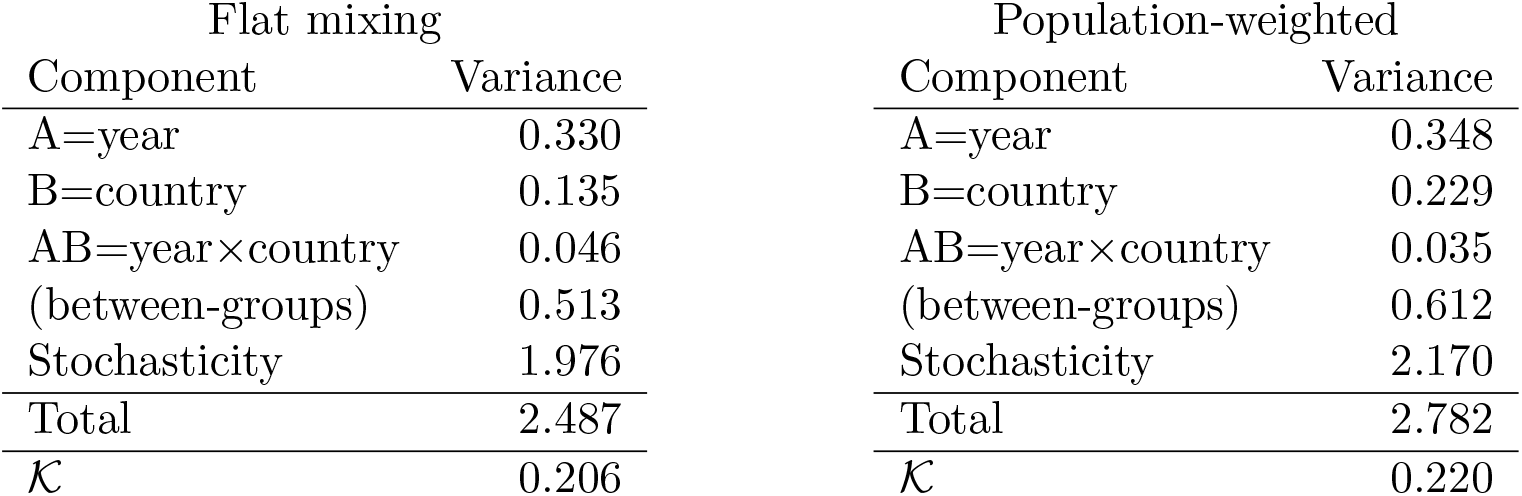
Components of variance in lifetime reproduction as affected by time (1960 compared with 2000) and country. Based on data from van Daalen and Caswell (2015).

Using the population-weighted rank-one mixing distribution makes only small changes in the variance components. The component due to years is similar to that with a flat mixing distribution. The component due to countries is larger than that in the flat mixing distribution. This contrasts with the results for longevity in Table 1, when weighting the races by population size reduced, not increased, the variance due to race. The interaction is again very small, and the variance ratio 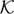 = 0.22 is very similar to that with the flat mixing distribution.

#### 5.2.2 Variance in reproduction due to diet and pesticide exposure

There is a rich biodemographic literature that applies demographic methods to laboratory populations of animals to evaluate the population effects of exposure to toxic substances.

We present one such example here, to demonstrate how exposure to conditions, that are known to affect human fertility, can be analysed in a case where experimentation is real, not imaginary.

Rotifers are microscopic invertebrate animals commonly used as bioindicator species for water quality, and increasingly used as model organisms for the study of ageing, maternal effects, and maternal investment (Bock et al., 2019). They have a relatively short lifespan (≈ 2 weeks), but high maternal investment into a small number of offspring.

In a classic experiment by Rao and Sarma (1986), the rotifer *Brachionus patulus* was exposed to five different concentrations of the pesticide DDT under low and high food resource levels. Age-specific survival and fertility schedules were reported for all combinations of DDT treatment and food level. The resulting arrays of means and variances of LRO are

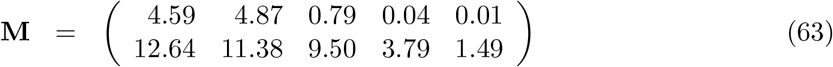

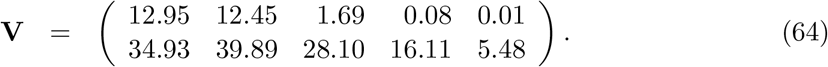

The two rows represent low and high food levels, respectively. The columns represent the five increasing levels of DDT exposure (0, 15, 30, 45, 60 *μ*g l^*−*1^ of DDT).

In this experiment, both DDT exposure and low food levels reduce mean lifetime reproduction. The effect of the pesticide exposure is greater at low food than at high food levels. The variance in LRO decreases with increasing DDT exposure and at low food levels. The variance is much larger than the mean, implying that the distribution of LRO is overdispersed relative to the Poisson distribution. This is a frequent pattern in analyses of LRO; it results from some portion of the population failing to reproduce at all (e.g., Tuljapurkar et al., 2020; van Daalen and Caswell, 2017).

The uniform mixing distribution is appropriate to a designed experiment like this. The resulting variance decomposition is shown in Table 4. The biggest component is that due to pesticide exposure, with food level contribution slightly less. The between-group variance, including the two main effects and their interaction, accounts for 57% of the variance in the experiment, which is higher than any of the percentages found to date for variance in longevity. The contribution of the food×DDT interaction is small, making up 9% of the total between-group variance.

**Table 4:**
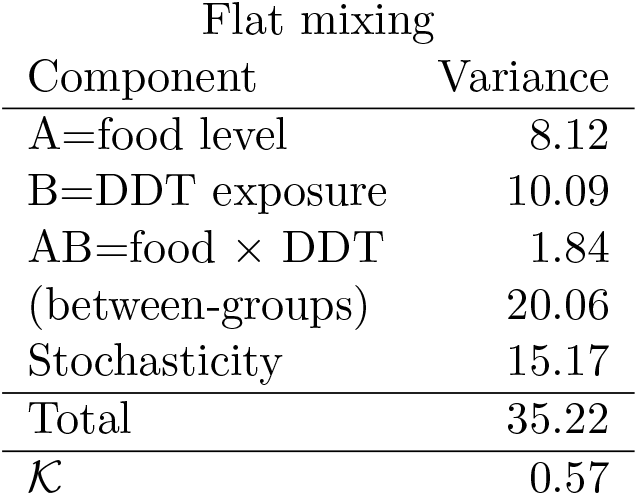
Components of variance in lifetime reproduction as affected by food level and DDT exposure. Data from the experimental study of Rao and Sarma (1986).

## 6 Discussion

Despite the obvious importance of variation, demography has often focused on expected values. Life expectancy is the expected value of longevity, the net reproductive rate R_0_ is the expected value of lifetime reproduction, and the total fertility rate TFR is the expected value of lifetime reproduction conditional on survival to the end of reproduction. The focus is expanding as questions of inequality become more important. Addressing these questions leads naturally to partitioning variation into components due to differences among individuals in the rates to which they are subject (heterogeneity) and components due to the stochastic outcomes of those rates among individuals subject to the same rates.

The current state of the art focuses on contributions of factors treated one at a time (sex, education, income, nutrition, etc.). In many studies of variance in longevity, it has been found that the contribution of heterogeneity, even in factors known to have important effects on individuals, is dwarfed by the contribution of stochasticity. Variance in lifetime reproduction has been studied less, but it appears that heterogeneity may make larger contributions to this demographic outcome.

Variance decomposition requires the means and variances of the outcome for all combinations of the factors. Although the means and variances can be obtained in many ways, Markov chains with rewards are a particularly powerful method. They can be applied to lifetime reproduction, to longevity in single-or multi-state models, and to healthy longevity for both prevalence-based (e.g., Caswell and Zarulli 2018 and Zarulli and Caswell 2022 for disability, Caswell and van Daalen 2021 for stages of cancer, Owoeye et al. 2020 for malnutrition), and incidence-based (Caswell and van Daalen 2021 for stages of cancer) models.

In this paper we have provided the results needed to apply these means and variances in studies of multiple factors operating simultaneously. The protocol for the variance decomposition follows a simple series of steps, independent of the number of factors considered.

### 6.1 A protocol for factorial variance decomposition

1. Define the factors that characterise the heterogeneity among individuals (e.g., sexes, races, income, education, resource levels, etc.) and the levels of each factor.
2. Choose a demographic outcome of interest (e.g., longevity, lifetime reproductive output).
3. Compute the means and variances of this demographic outcome for all combinations of the factors, using Markov chains, Markov chains with rewards, or life tables as appropriate.
4. Create the arrays **M** and **V** containing the means and variances, as in equations (13) and (14). The dimension of these arrays is the number of factors in the study (i.e., for *n* factors, **M** and **V** are n-dimensional).
5. Think about the question of interest and specify a set of individuals over which the variances are to be computed:
  a. a set consisting of equal representation of all factor combinations (the flat mixing distribution), or
  b. a set defined by a rank-one combination of marginal distributions.
6. Create the array **Π**, containing the *n*-dimensional mixing distribution, as in (15).
7. Treating all the factor combinations as groups, calculate the overall within- and between-factor variance components *V*_within_ and *V*_between_ using equations (9) and (10).
8. Partition the between-group variance into components:
  a. compute marginal mean arrays for each factor and each factor interaction, as in equations (20)–(22),
  b. create the corresponding marginal mixing distributions, as in equations (23)–(25), and
  c. compute the variance components for each factor and each interaction, as in equations (27)–(29).

### 6.2 Variance as a measure of inequality

There are many ways to measure the variation (‘inequality’ in a broad sense) of some quantity. Economists have developed many indices to address specifically economic issues related to income, transfers, and so on (e.g., Jenkins and van Kerm, 2009; Atkinson, 2015). In demographic contexts, these measures are highly correlated among each other (Van Raalte and Caswell, 2013), so if interests focuses only on the value of the index, there is little to choose among them. But to go beyond the values, there are properties of the variance that make it an attractive choice.

As is well known, the variance is decomposable into within- and between-group components (as are indices based on entropy). Additive decomposability is more than a mathematical nicety; it is fundamental to our attempts to understand how heterogeneity contributes to inequality of outcomes. It was for this purpose that Fisher introduced it as the basis for the analysis of experimental data (Fisher, 1936). Statistical ANOVA has developed into an enormous variety of study designs, referred to as ‘experimental designs’ in the experimental sciences, and that correspond in our context to arrangements of factors within and across populations. The factorial study design we explore here only scratches the surface of the possibilities.

The most popular alternative to the variance is the Gini coefficient. The Gini coefficient is based on the mean of the absolute values of the deviations between pairs of values (e.g., individuals) selected from the distribution. Because of this, Permanyer et al. (2023) call the Gini coefficient an individual measure. They contrast it with the variance, which they call a group measure, because it is usually written as the mean of the squared deviations from the mean, rather than among values. However, this distinction does not hold up on closer examination. The mean difference is

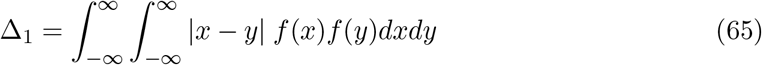

The Gini coefficient is a standardised, dimensionless version of the mean difference

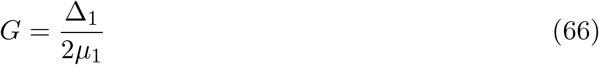

The variance can also be written as a difference among individuals, “without reference to deviations from a central value, the mean” (Kendall and Stuart, 1969, p. 47) as

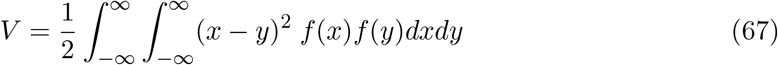

The similarity between the mean difference and the variance is clear; the former takes absolute values of the differences between individuals, the latter the square of the differences. Just as the Gini coefficient is a standardisation of the mean difference, the variance can be non-dimensionalized in the form of the familiar coefficient of variation

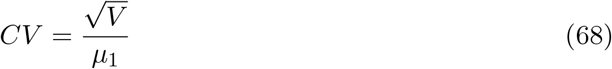

or the standardised variance

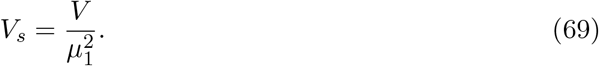

The standardised variance is known in evolution and anthropology as Crow’s index of the opportunity for selection (Crow, 1958; van Daalen and Caswell, 2024; Courtiol et al., 2012). It can be additively decomposed into between-group and within-group components as a measure of inequality (Rosenbluth, 1951).

The variance also has the advantage of being a moment of the distribution, inviting connections to other moments, which are easily calculated by Markov chain methods. These moments highlight other aspects of the distribution (e.g., skewness), and at least some of them can also be decomposed into within- and between-group components. There seems to be no comparable linkage of absolute deviations to such other properties of the distribution.

### 6.3 Remarks

The variance partitioning presented here is related to, but not quite the same as, statistical Analysis of Variance (ANOVA). Our results do not provide hypothesis tests, because they lack an underlying sampling theory to obtain distributions of the variance components under a null hypothesis. If they were, the between-group components would usually be very insignificant. However, the factors investigated in demographic studies are usually known a priori to be of social or biological importance. That the contribution of heterogeneity to variance is small does not imply that the factors are not worthy of attention; it implies only that stochasticity is itself an important factor worthy of study (see Caswell 2023, Section 8.5).

A recent survey of single-factor studies found that heterogeneity usually explained only 5%–10% of the variance in longevity. The results examined here suggest that including multiple factors and their interactions may not increase K very much. Further comparative research is needed. The variance ratio for lifetime reproduction appears to be larger than that for longevity, and the variance components appear to be strongly influenced by conditions. van Daalen et al. (2022) found that heterogeneity in maternal age in a rotifer explained about 26% of the variance in lifetime reproduction under laboratory conditions, but as little as 2% under most conditions that would lead to a stationary population. The sensitivity analysis of variance components (van Daalen and Caswell, 2020) may help explore these relationships.

We encourage the use of the methods we present here to explore the contributions of multiple factors and their interactions to variance in demographic outcomes. Patterns await to be discovered.

## 7 Acknowledgements

This research has been supported by the European Research Council under ERC Advanced Grant 322989 (INDSTOCH) and ERC Advanced Grant 788195 (FORMKIN). S.F.v.D. was also supported by the Postdoctoral Scholar Program at Woods Hole Oceanographic Institution, with funding provided by the Doherty Foundation. HC acknowledges with gratitude the late Dr. John L. Gill of Michigan State University.

## Footnotes

1 It is convenient to think of a a finite number of groups, but the concepts and theory are essentially the same for continuous heterogeneity. A familiar example is the gamma-Gompertz distribution of mortality, which can be discretised to allow variances to be calculated (Caswell, 2014).

2 An exception to this generalisation is studies in which groups are defined as quantiles of the distribution of some variable in a population, as in distributions of income, neighbourhood deprivation indices, etc.

3 An alternative approach, using small variance approximations, has been given by Snyder and Ellner (2018, 2016). Their conclusions are the same as ours.

4 The countries are Austria, Bulgaria, Canada, Switzerland, Czech Republic, East Germany, West Germany, Estonia, Finland, France, Scotland, England and Wales, Hungary, Japan, Lithuania, Netherlands, Portugal, Russia, Slovakia, Sweden, Ukraine, United States, Australia, Belgium, Belarus, Denmark, Spain, Ireland, Italy, New Zealand, and Poland.

## Notes

### Competing Interest Statement

The authors have declared no competing interest.

